# Senescence-associated KRAS upregulation in peripheral T cells links to premature coronary artery disease

**DOI:** 10.64898/2026.09.22.753659

**Authors:** Zhao Sun, Ziyu Yang, Xuedong Wang, Meng Zhang, Rui Gao, Mengyue Yang, Yanchao Li, Qi Liu, Jingbo Hou

## Abstract

**Aims:** Premature coronary artery disease (PCAD) lacks specific molecular drivers, and the role of immunosenescence is unclear. We investigated whether aging-related gene dysregulation in T cells contributes to PCAD.

**Methods:** We combined bulk transcriptomics of PBMCs from 12 PCAD patients and 21 controls, single-cell RNA sequencing of PBMCs and human atherosclerotic plaques, weighted gene co-expression network analysis, gene perturbation network analysis, and molecular docking.

**Results:** KRAS was identified as a hub gene intersecting PCAD-associated genes and aging-related genes. Single-cell analysis showed KRAS upregulation predominantly in effector CD8+ T cells, which exhibited the highest senescence scores that were further elevated in disease. Network perturbation of KRAS strongly impacted the cell killing pathway. KRAS-high effector CD8+ T cells were detected in coronary and carotid plaques, displaying enhanced cytotoxicity, exhaustion, and senescence features. Additionally, a candidate small molecule was computationally predicted to bind inactive KRAS.

**Conclusions:** Elevated KRAS expression in senescent, cytotoxic CD8+ T cells is associated with PCAD, bridging immunosenescence and premature atherosclerosis. This finding provides a novel biomarker candidate and potential therapeutic entry point, awaiting further functional validation.

## INTRODUCTION

PCAD is an increasingly serious and distinctively challenging subset of cardiovascular disease^[1]^. For decades, it attracted relatively little attention, largely because the incidence of coronary disease in young adults was substantially lower than in older populations^[2]^. Yet cardiovascular mortality has remained a leading cause of death among the young, and the proportion of AMI occurring in younger patients has risen steadily over time. Current primary prevention strategies rely predominantly on risk scores derived from middle-aged and elderly cohorts, which may systematically underestimate risk in younger individuals. As a result, PCAD is often diagnosed only after an acute cardiovascular event, and the development of more precise preventive and therapeutic strategies is constrained by a lack of robust epidemiological evidence^[3, 4]^. The risk factors and subclassification of PCAD also remain poorly defined. Although a majority of young patients present with traditional risk factors such as smoking, obesity, and hyperlipidemia, the accelerated course of atherosclerosis and its premature onset warrant further investigation^[5–7]^. Collectively, these clinical challenges underscore the urgent need to identify specific molecular drivers of PCAD—drivers that likely extend beyond the conventional cardiometabolic pathways.

Aging is a well-established independent risk factor for CAD^[8]^. With advancing age, both innate and adaptive immunity decline, a process collectively termed immunosenescence. This functional deterioration is characterized by dysregulated cytokine secretion, impaired intercellular communication, and altered proportions of immune cell populations. Immunosenescence compromises the capacity to eliminate pathogens and senescent cells and disrupts the normal resolution of inflammatory responses, ultimately fostering a state of chronic inflammation that has been shown to promote atherosclerosis^[9]^. As central players in adaptive immunity, senescent T cells accumulate within atherosclerotic plaques of older patients with coronary artery disease, where they exhibit a diminished ability to process oxidized low-density lipoprotein antigens and secrete large quantities of chemokines that exacerbate inflammation in plaques^[10, 11]^. In the context of PCAD, however, the contribution of T-cell immunosenescence remains poorly understood.

A key member of the RAS proto-oncogene family, KRAS protein encodes a small GTPase that regulates cell proliferation, differentiation, and survival through signaling cascades such as MAPK/ERK, PI3K/AKT/mTOR, Ral GEF/Ral A/B, JAK/STAT3, and NF-κB^[12–14]^.Abnormal KRAS activation—whether driven by mutation or wild-type overexpression—predominantly triggers oncogene-induced cellular senescence rather than malignant proliferation in the absence of cooperating genetic events^[15]^, revealing an intrinsic link between KRAS signaling and senescence regulation. Indeed, wild-type KRAS overexpression has been shown to impair the effector function of CD8+T cells via the ERK pathway^[16]^. In the context of atherosclerosis, copy number gains at the KRAS locus have been detected in human carotid plaque specimens, suggesting its involvement in vascular pathology^[17]^. However, a critical question remains unexplored: in PCAD, whether dysregulated KRAS expression in T cells—rather than oncogenic mutations—drive accelerated immunosenescence and tumor-like immune effector programs, thereby promoting arterial injury.

In this study, we integrated transcriptomic profiling, scRNA-seq, virtual gene knockout, deep learning-based compound prediction, and molecular docking to characterize the role of KRAS in peripheral T cells of patients with PCAD. We identified KRAS as a hub gene at the intersection of PCAD-associated expression signatures and aging-related gene sets, with its upregulation predominantly localized to effector CD8+T cells. Single-cell transcriptomic analysis coupled with an AI-assisted virtual knockout algorithm revealed that elevated KRAS drives a cytotoxic effector program and is closely associated with features of cellular senescence. We further confirmed the presence of this KRAS-high cytotoxic effector CD8+T-cell subpopulation in single-cell transcriptomic data from human coronary atherosclerotic plaques and in carotid atherosclerotic lesions. Moreover, using a transcriptome-driven artificial intelligence model, we predicted a novel small-molecule candidate capable of binding KRAS, offering a potential strategy for targeting a protein traditionally regarded as undruggable^[14]^. Taken together, our findings identify KRAS upregulation in T cells as a novel nexus linking accelerated immunosenescence to premature atherosclerosis, providing both mechanistic insights and a candidate therapeutic direction for this understudied patient population. The workflow of this study is illustrated in Figure 1.

**Figure 1.**
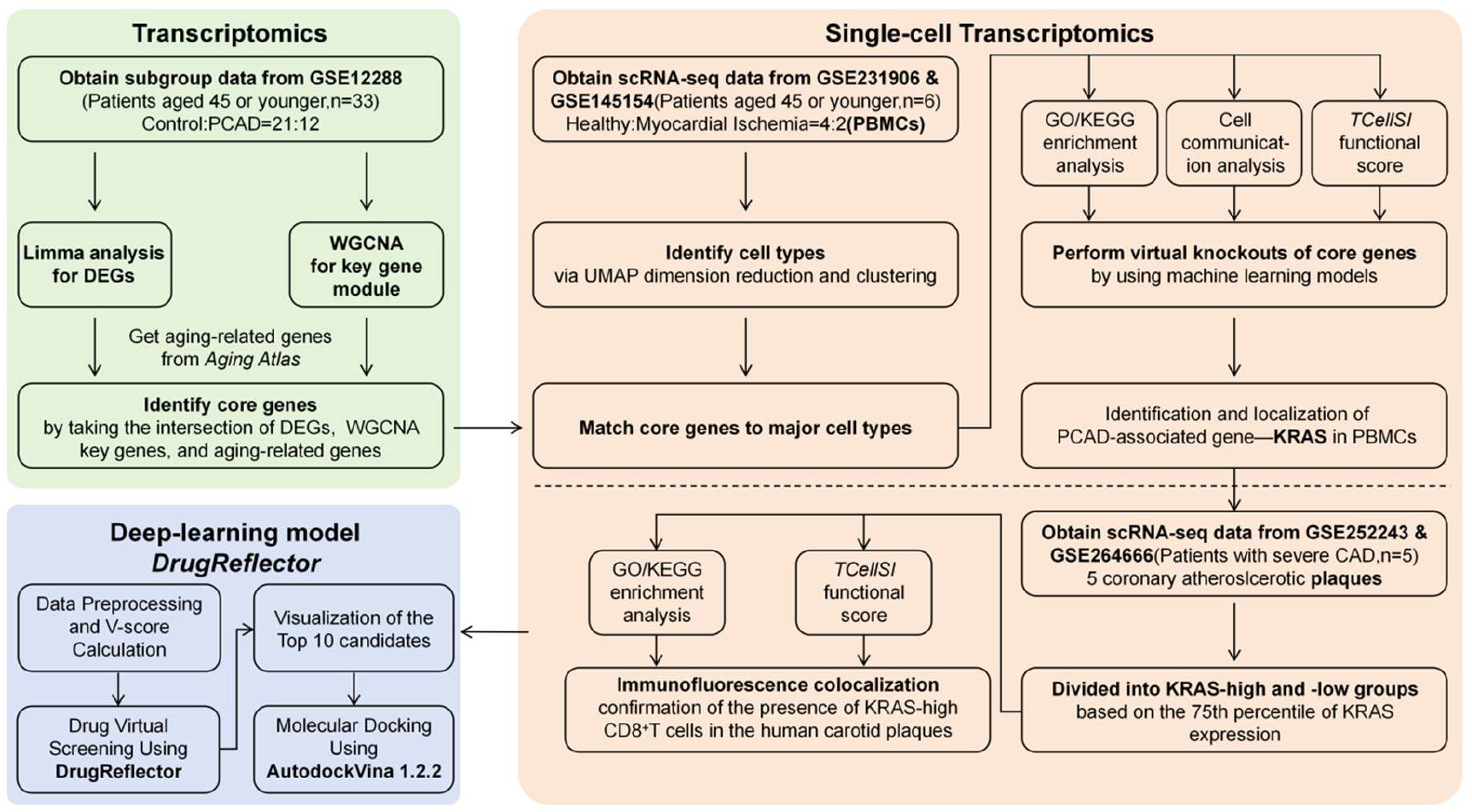
Workflow of this study.

## MATERIAL AND METHODS

### Public Data Acquisition

The bulk RNA-seq dataset of peripheral blood mononuclear cells (PBMCs) from patients with premature coronary artery disease and healthy controls was obtained from the Gene Expression Omnibus (GEO) under accession number GSE12288. Single-cell RNA-seq (scRNA-seq) data of PBMCs were retrieved from GEO (GSE231906 and GSE145154). scRNA-seq data of human coronary atherosclerotic plaques were downloaded from GEO (GSE252243 and GSE264666). Detailed sample information and inclusion criteria are provided in the original publications.

### Clinical Sample Collection and Validation

We conducted a case-control validation study using clinical samples collected at The Second Affiliated Hospital of Harbin Medical University between January 1, 2025, and January 1, 2026. The study was approved by the Ethics Committee of The Second Affiliated Hospital of Harbin Medical University (approval number: KY-2025394), and written informed consent was obtained from all participants or their legal representatives.

For PBMC validation, we enrolled 5 patients with premature coronary artery disease (PCAD) and 5 control subjects. All participants were younger than 45 years and underwent coronary angiography. PCAD cases were defined as angiographic coronary stenosis >70% in at least one major epicardial vessel. Controls were defined as individuals with no detectable coronary stenosis on angiography. Cases and controls were not matched. PBMCs were isolated from peripheral blood, and mRNA levels of KRAS, SYK, and TAB2 were measured by quantitative real-time PCR (qPCR) as described below.

For plaque validation, formalin-fixed paraffin-embedded carotid atherosclerotic plaque specimens were obtained from 5 patients with symptomatic severe carotid stenosis who underwent carotid endarterectomy (CEA). The diagnosis was confirmed by digital subtraction angiography (DSA) and met at least one of the following criteria: (1) transient ischemic attack (TIA) or minor non-disabling stroke within 6 months, with ipsilateral carotid stenosis ≥50%; or (2) stenosis between 50% and 69% with unstable plaque features (e.g., surface ulceration, hemorrhage, or echogenic plaques with active embolic risk) and an assessed risk of recurrent stroke.

Clinical data including age, sex, smoking status, hypertension, diabetes, and lipid profiles were collected. These potential confounders were adjusted for in the statistical analysis. All experimental procedures and result interpretations were performed under double-blind conditions.

### Differential Expression Analysis and Weighted Gene Co-expression Network Analysis (WGCNA)

For the bulk transcriptomic data, raw expression values were normalized and differentially expressed genes (DEGs) between CAD and control groups were identified using the limma package (version 3.62.2) in R (version 4.4.3). Genes with a p-value < 0.05 and an absolute fold change > 1.5 were considered significant.

Weighted gene co-expression network analysis was performed using the WGCNA package (version 1.74). A signed network was constructed with a soft-thresholding power of 12, selected to achieve approximate scale-free topology. Modules were identified using dynamic tree cutting with a minimum module size of 80. The module eigengenes were correlated with clinical traits to identify the module most strongly associated with CAD. The intersection of module key genes, DEGs, and aging-related genes from the Aging Atlas database (https://ngdc.cncb.ac.cn/aging/index) was visualized with a Venn diagram.

### Single-Cell RNA-Seq Data Processing

PBMC and plaque scRNA-seq data were processed using the Seurat package (version 5.5.0). For each dataset, cells with < 200 detected genes and > 10% mitochondrial reads were filtered out. Data were normalized using the SCTransform method. Principal component analysis (PCA) was performed on the top 2000 highly variable genes, and the first 20 principal components were used for UMAP dimensionality reduction and graph-based clustering (resolution = 0.6). Cell types were annotated based on canonical marker genes: T cells (CD3D,CD3E), NK cells (KLRD1,GNLY), B cells (MS4A1,CD79B), monocytes (CD14,LYZ), dendritic cells (CDKN1C,LILRB2), macrophages (CD68,C1QA), smooth muscle cells (ACTA2,MYH11), fibroblasts (COL6A1,FN1), and endothelial cells (PECAM1,EGFL7).

### Cell-Cell Communication and Pathway Analysis

Intercellular communication was inferred using CellChat (version 2.1.1) with the default ligand-receptor database. The number and strength of inferred interactions were compared between groups. Gene set variation analysis (GSVA) was performed using the GSVA package (version 2.0.7) with hallmark gene sets from the Molecular Signatures Database (MSigDB). For T-cell-specific differential expression, DEGs between the MI and Healthy groups were identified using the FindMarkers function (log-fold change threshold > 0.5, adjusted p < 0.05) and subjected to GO and KEGG enrichment analysis using clusterProfiler (version 4.14.6).

### T-Cell Subset Analysis and Functional State Scoring

T cells were re-clustered and visualized with UMAP. Subsets were defined based on the expression of established markers: naive CD4+T cells (CD4,CCR7,SELL), memory CD4+T cells (CD4,CXCR4, RORA), CD4+ regulatory T cells (CD4,FOXP3, IL2RA), naive CD8+ T cells (CD8A,CCR7, SELL), and effector CD8+ T cells (CD8A,GZMK,GZMB). Functional state scores (cytotoxicity, exhaustion, senescence) were calculated for each cell using the TCellSI package (version 1.2.0) with default parameters.

### Virtual Gene Knockout Analysis

The scTenifoldKnk package (version 1.0.3) was used to perform in silico knockout of KRAS in T cells. The algorithm constructs a single-cell-level gene regulatory network, removes the target gene, and compares the resulting network to the original to identify significantly perturbed genes. The top 20 most perturbed genes were retained for GO and KEGG enrichment analysis.

### Deep Learning-Based Drug Prediction and Molecular Docking

The deep learning model DrugReflector was trained on the Connectivity Map (CMap) database. The list of upregulated DEGs from MI vs. Healthy T cells was used as the input disease signature. The model returned ranked compounds based on their predicted ability to reverse the signature. The top-ranked compound BRD-K78432605 was selected for molecular docking. The crystal structure of the GDP-bound inactive KRAS protein (PDB: 4EPR, resolution 2.00 Å) was retrieved from the Protein Data Bank. Molecular docking was performed using AutoDock Vina (version 4.2.6) with a grid box centered at the SOS-binding interface. The docking result was visualized with PyMOL (version 2.6.0).

### Quantitative Real-Time PCR

Tissue was ground into powder and added to TRIzol Reagent (Invitrogen, USA), while cell samples were added to TRIzol Reagent after discarding the supernatant. The procedure of RNA extraction was as follows: chloroform (350 μL per 1 mL TRIzol;Xilong Scientific) was added, and the mixture was then centrifuged for 15 min at 12000 rpm at room temperature. The supernatant was mixed with 500 μL of isopropanol (Xilong Scientific) and left to stand for 10 min at room temperature, then centrifuged for 10 min at 12000 rpm at 4◦C. After discarding the supernatant, 1 mL of 75% anhydrous ethanol (Xilong Scientific) was added and centrifuged for 5 min at 7500 rpm at 4◦C. The supernatant was discarded, and the white solid at the bottom was preserved; when the precipitate was sufficiently dried, enzyme-free water was added, and the concentration of RNA was measured and the RNA was stored at − 80^◦^C for later use. Subsequently, cDNA was obtained by reverse transcription reaction using Toyobo reverse transcription kit, followed by qRT-PCR reaction using 2X M5 HiPer SYBR Premix EsTaq plus (with Tli RNase H) (Mei5bio).Relative gene expression was calculated using the 2 ^-ΔΔCt method. Data were normalized to GAPDH for mRNA targets The primer sequences for the target gene were as follows:

TAB2:

Forward: 5’-GCCACCAAATTGATTTTCAGGTT-3’.

Reverse: 5’-TGCGTAGACCAGAAATTCCAGA-3’. SYK:

Forward: 5’-CATGGAAAAATCTCTCGGGAAGA-3’.

Reverse: 5’-GTCGATGCGATAGTGCAGCA-3’.

KRAS:

Forward: 5’-ACAGAGAGTGGAGGATGCTTT-3’.

Reverse: 5’-TTTCACACAGCCAGGAGTCTT-3’. GAPDH:

Forward, 5’-GAGAAGGCTGGGGCTCATTT-3’.

Reverse, 5’-ATGACGAACATGGGGGCATC-3’.

### Immunofluorescence Staining of Atherosclerotic Plaques

Formalin-fixed paraffin-embedded human carotid atherosclerotic plaque specimens were sectioned at 4 μm thickness. Serial sections were deparaffinized, rehydrated, and subjected to heat-induced antigen retrieval in citrate buffer (pH 6.0) using a microwave. After blocking with 5% bovine serum albumin for 1 h at room temperature, sections were incubated overnight at 4 °C with primary antibody pairs as follows:

1. anti-CD8a(mouse,1:200,Proteintech,Cat No.66868-1-Ig)+ anti-KRAS(rabbit,1:200,Proteintech,Cat No.12063-1-AP);
2. anti-KRAS(rabbit)+anti-p16-INK4A(mouse,1:200,Proteintech,Cat No.60626-1-Ig).

After washing, sections were incubated with Alexa Fluor 488-conjugated goat anti-rabbit IgG and Alexa Fluor 594-conjugated goat anti-mouse IgG (1:500, Invitrogen) for 1 h at room temperature. Nuclei were counterstained with DAPI.

Images were captured using a Zeiss Axio Observer fluorescence microscope and analyzed with ImageJ software. For each staining pair, five random high-power fields (×200) were selected, and double-positive cells were counted manually.

### Statistical Analysis

All statistical analyses were performed in R (version 4.4.3). For clinical validation, comparisons between PCAD cases and controls were performed using Student’s t-test for continuous variables, with adjustment for potential confounders including age, sex, smoking, hypertension, diabetes, and lipid profiles using multivariable linear regression. Multiple-group comparisons were conducted with one-way ANOVA followed by Tukey’s post-hoc test. A p-value < 0.05 was considered statistically significant, unless otherwise specified. For the carotid plaque immunofluorescence validation, five random high-power fields (×200) per staining pair were selected, and double-positive cells were counted manually; no formal statistical testing was performed for this descriptive validation.

## RESULTS

### Identification of aging-related hub genes in PBMCs of patients with PCAD

First,we retrieved PBMCs transcriptomic data from public databases(GSE12288) for 12 coronary artery disease (CAD) patients aged <45 years and 21 healthy controls. After regrouping samples into PCAD and Control groups, limma differential expression analysis with thresholds of p < 0.05 and |fold change| > 1.5 identified 2,289 differentially expressed genes (DEGs), of which 1,175 were upregulated and 1,114 downregulated (Figure 2A). WGCNA on the full gene set under the same grouping yielded 23 modules; among them, the purple module showed the strongest association with PCAD (Figure 2B). Intersecting the 315 key genes from this module with the 2,289 DEGs and 503 aging-related genes from the Aging Atlas database revealed three core regulatory genes: TAB2, SYK, and KRAS (Figure 2C). All three were expressed at higher levels in PBMCs of the PCAD group than in the Control group (Figure 2D), and these results were further validated by qPCR (Figure 2E).

**Figure 2.**
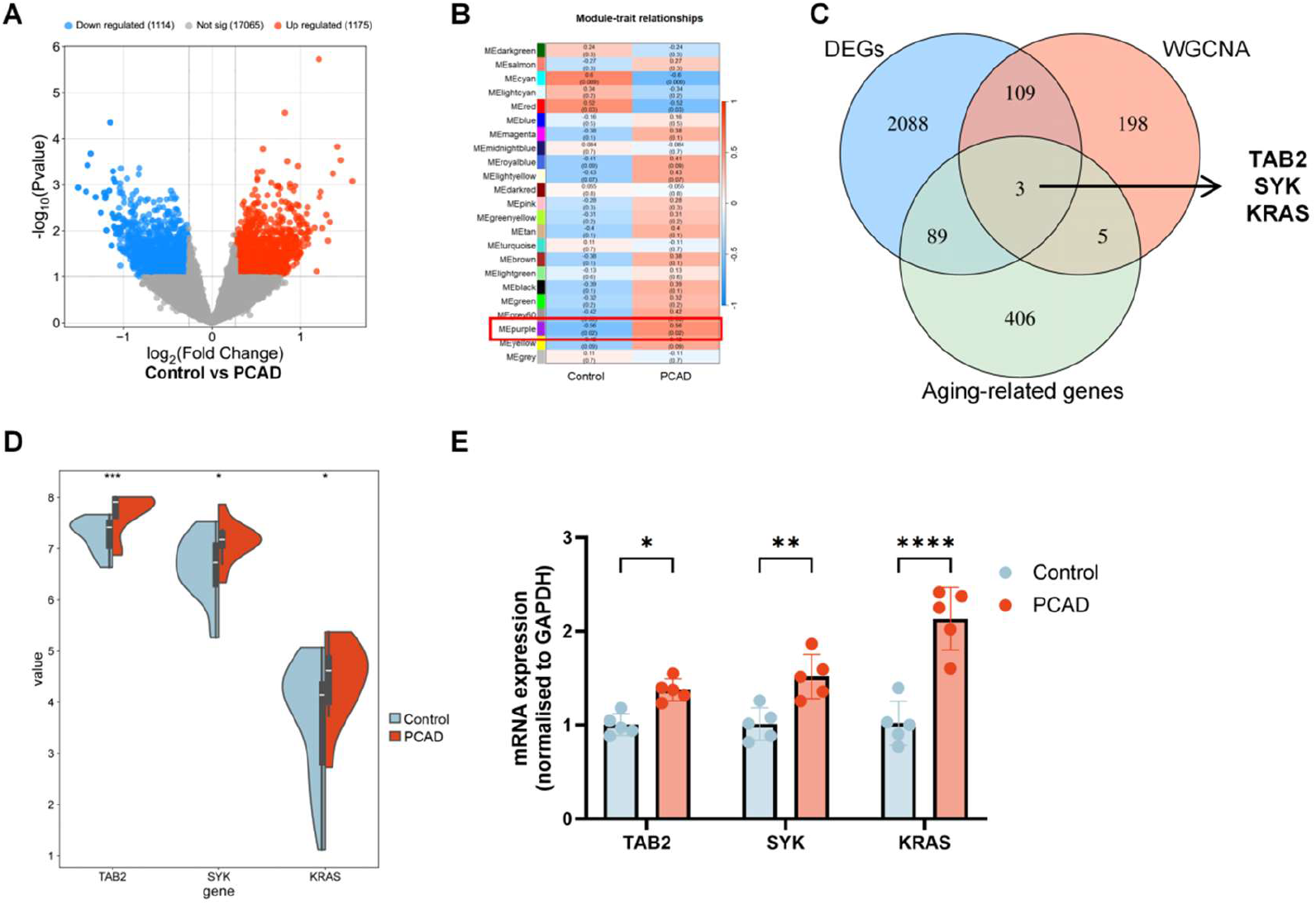
Identification of aging-related hub genes in PBMCs of patients with premature CAD. (A) Volcano plot of differentially expressed genes (DEGs) between CAD patients and healthy controls. (B) Heatmap of module–trait associations derived from weighted gene co-expression network analysis (WGCNA); the purple module shows the strongest correlation with CAD. (C) Venn diagram illustrating the overlap among 315 key genes from the purple module, 2,289 DEGs, and 503 aging-related genes from the Aging Atlas database, yielding three shared hub genes: TAB2, SYK, and KRAS. (D) Violin plots depicting the expression levels of TAB2, SYK, and KRAS in PBMCs from the CAD group versus the Control group. (E) qPCR validation of the expression differences of the three hub genes. Data are presented as mean ± SD; *p* < 0.05, **p* < 0.01, ***p* < 0.001.

### Upregulation of KRAS in peripheral blood T cells is closely linked to PCAD

To delineate the cellular context and functional relevance of the identified core genes in premature CAD, we re-analyzed publicly available scRNA-seq data (GSE231906 & GSE145154) of PBMCs from 2 patients with ischemic cardiomyopathy (myocardial ischemia group, MI group) aged <45 years and 4 healthy controls (Healthy group) without coronary lesions. After dimensionality reduction, clustering, and annotation, five major cell types were identified: T cells, NK cells, B cells, monocytes, and dendritic cells. Compared with the Healthy group, the MI group exhibited a higher proportion of T cells and a markedly lower proportion of monocytes (Figure 3A and B). To determine the distribution of TAB2, SYK, and KRAS across cell types, we calculated their expression levels in each cell type across groups. TAB2 and KRAS were predominantly expressed in T cells, whereas SYK was mainly expressed in monocytes (Figure 3C), and all three genes showed elevated expression across cell types in the MI group relative to the Healthy group (Figure 3D). Cell-cell communication analysis revealed that T cells occupied a central hub position among all cell types (Figure 3E). Notably, both the number and strength of inferred interactions in PBMCs were globally reduced in the MI group compared with the Healthy group (Figure 3F). Although pathways such as CypA and MIF remained dominant in the intercellular communication network of the MI group, their overall signaling activity was still markedly weaker than that in the Healthy group (Figure 3G). Gene set variation analysis (GSVA) across all PBMCs showed that the KRAS signaling pathway was significantly activated in the MI group, and the aging-associated p53 pathway also exhibited relatively high activity (Figure 3H).

**Figure 3.**
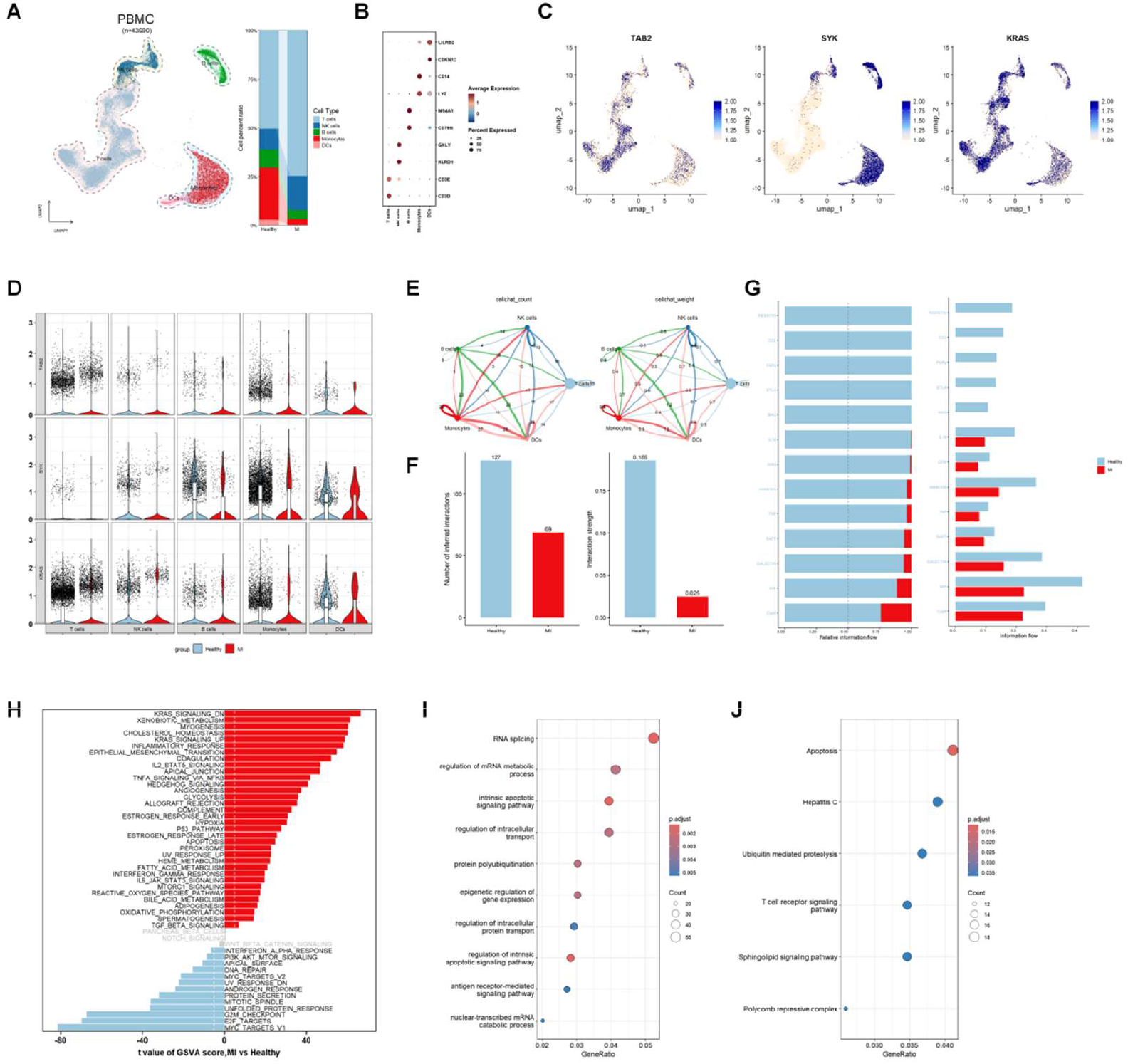
PBMC single-cell sequencing results display the core genes. UMAP plot of PBMC and cell type proportions; (B) Bubble plot of marker gene expression in different cell types; (C) Expression distribution of TAB2, SYK, and KRAS; (D) Expression levels of TAB2, SYK, and KRAS in different groups and cell types (shown as violin plots); (E) CellChat cell communication results (number and weight); (F) Comparison of CellChat cell communication results between different groups (number and weight); (G) Activity levels of interaction pathways in different groups; (H) Heatmap of GSVA pathway activity scores; (I) Bubble plot of GO enrichment analysis of upregulated genes in T cells of the MI group; (J) Bubble plot of KEGG enrichment analysis of upregulated genes in T cells of the MI group.

Taken together, these observations indicate that T cells occupy a central position in the PBMCs of the MI group: their proportion is significantly elevated, they exhibit the highest communication strength and interaction counts, and they serve as the principal cell type in which KRAS and its downstream signaling operate. T cells are therefore likely to play a key regulatory role in premature CAD. We subsequently performed differential gene expression analysis between T cells of the MI and Healthy groups, and subjected the genes upregulated in MI T cells to GO/KEGG enrichment analysis. GO analysis revealed significant enrichment of terms such as RNA splicing, mRNA metabolic process, and intrinsic apoptosis signaling pathway (Figure 3I), while KEGG analysis confirmed a prominent enrichment of the apoptosis pathway (Figure 3J). These results suggest that elevated KRAS expression in MI T cells is associated with enhanced apoptotic activity in peripheral blood T cells.

### KRAS-upregulated effector CD8+T cells undergo accelerated immunosenescence

To further clarify the precise localization and function of KRAS within the T-cell compartment, we performed subclustering and analysis of T cells. Based on marker gene expression, five T-cell subsets were identified: naive CD4+T cells, memory CD4+T cells, CD4+regulatory T cells, naive CD8+T cells, and effector CD8+T cells. In terms of proportions, the MI group exhibited a higher percentage of naive CD4+T cells and a relatively lower percentage of naive CD8+T cells, whereas the remaining three subsets showed minimal differences between groups (Figure 4A and B). KRAS was expressed across all T-cell subsets, with higher expression levels and detection rates observed in naive CD4+T cells and effector CD8+T cells (Figure 4C and D). We applied the TCellSI algorithm to score the functional states of different T-cell subsets across groups. Notably, the senescence score of effector CD8+T cells was the highest among all subsets and was further elevated in the MI group compared with the Healthy group, suggesting that KRAS-high effector CD8+T cells in the MI group may be undergoing accelerated immunosenescence.

**Figure 4.**
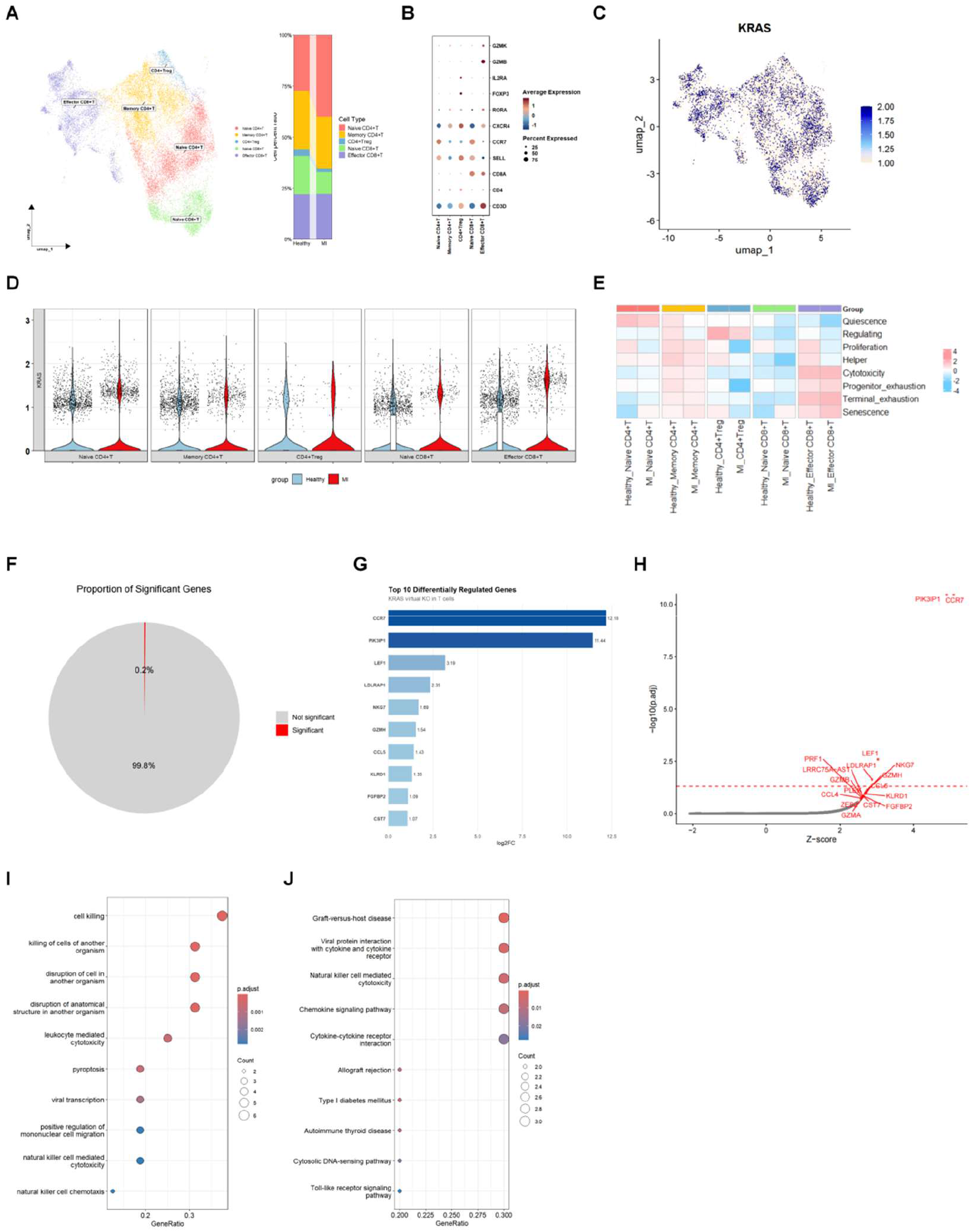
Functional analysis of T cell subsets in PBMC. A) UMAP plot of T cell subsets and cell type proportions; (B) Bubble plot of marker gene expression in different cell types; (C) Expression distribution of KRAS; (D) Expression level of KRAS in each T cell subset; (E) Heatmap of TCellSI functional scores; (F) Pie chart of the proportion of significantly perturbed genes after scTenifoldKnk virtual knockout; (G) Bar plot of the top 10 most significantly perturbed genes; (H) Scatter plot of significantly perturbed genes; (I) Bubble plot of GO enrichment analysis of significantly perturbed genes; (J) Bubble plot of KEGG pathway enrichment analysis of significantly perturbed genes.

To infer the core regulatory direction of KRAS from the single-cell transcriptomic network, we performed virtual knockout analysis of KRAS in T cells using scTenifoldKnk. The results showed that significantly perturbed genes accounted for approximately 0.2% of all genes, with CCR7 and PIK3IP1 being the most prominently affected (Figure 4F-H). GO and KEGG enrichment analyses of the top 20 perturbed genes revealed a striking enrichment of the cell killing pathway (Figure 4I and J), indicating a tight functional coupling between KRAS signaling and the cytotoxic effector program of T cells.

### Effector CD8+T cells with high KRAS expression infiltrate atherosclerotic plaques and exhibit enhanced effector function

Having established that KRAS upregulation in peripheral blood effector CD8+T cells enhances their cytotoxicity and promotes their senescence in patients with PCAD, we next examined whether this phenomenon also occurs within atherosclerotic plaques. We collected publicly available scRNA-seq data of coronary plaque samples from five patients with advanced coronary atherosclerosis and performed re-analysis(GSE252243 & GSE264666). After dimensionality reduction and clustering, seven cell types were annotated: macrophages, T cells, NK cells, B cells, smooth muscle cells, fibroblasts, and endothelial cells, all of which were distributed across the different plaque samples (Figure 5A and B). Consistent with the PBMC findings, KRAS exhibited the highest expression proportion and overall expression level in T cells (Figure 5C and D). Following subclustering of T cells using the same markers as those applied to PBMCs, four T-cell subsets were identified: naive CD4+T cells, memory CD4+T cells, CD4+regulatory T cells, and effector CD8+T cells (Figure 5E and F). To determine whether KRAS-high effector CD8+T cells in plaques share similar functional characteristics with their peripheral counterparts, we divided the effector CD8+T cells into KRAS-high and KRAS-low groups using the upper quartile (75th percentile) of KRAS expression as the cutoff.

**Figure 5.**
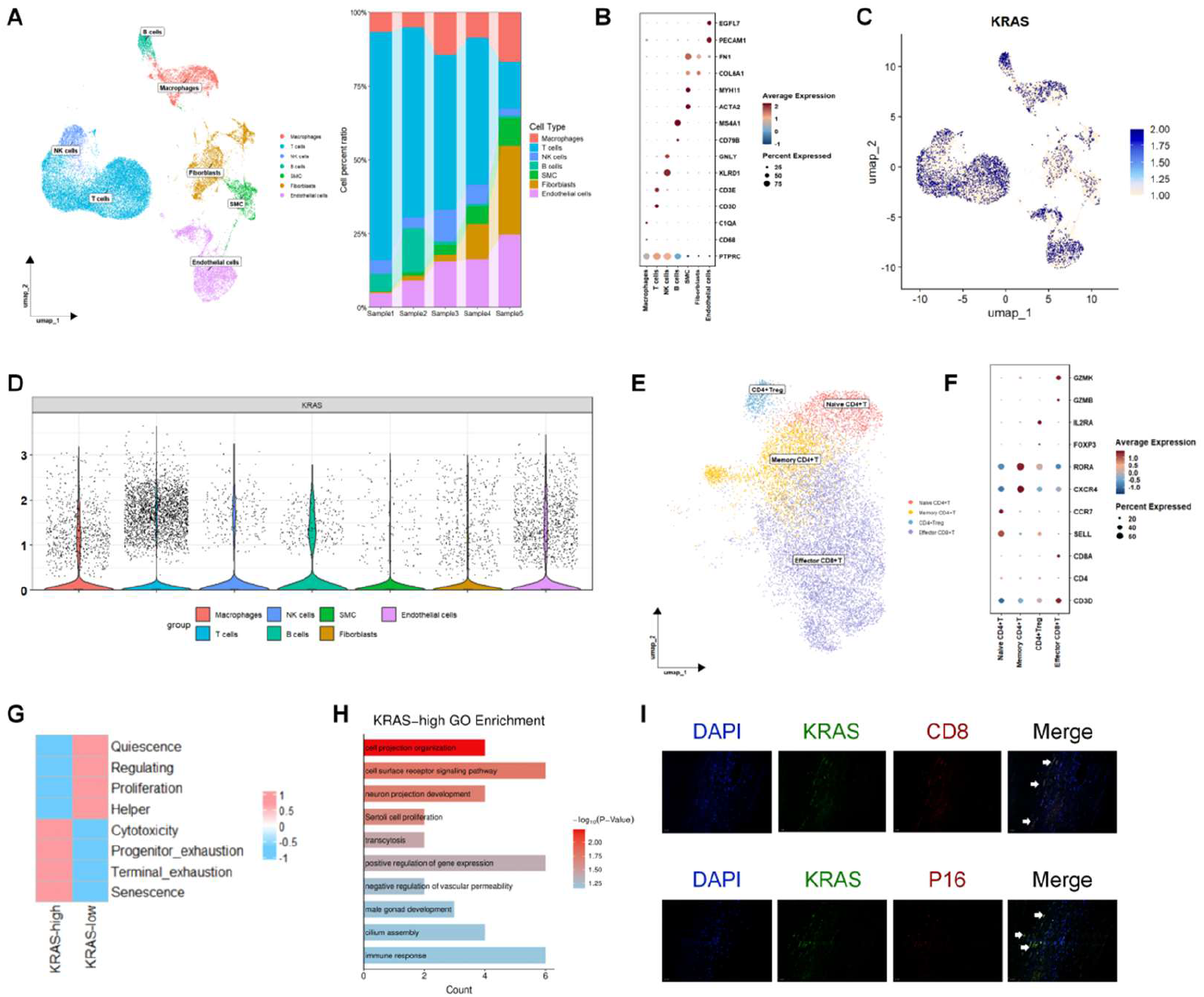
Single-cell sequencing results of coronary artery plaques verify the existence and function of the T cell subset with KRAS upregulation. (A) UMAP plot of coronary artery plaques and cell type proportions; (B) Bubble plot of marker gene expression in different cell types within plaques; (C) Expression distribution of KRAS; (D) Expression level of KRAS in different cell types; (E) UMAP plot of T cell subsets; (F) Bubble plot of marker gene expression in each T cell subset within plaques; (G) Comparison of TCellSI functional scores between KRAS-high and KRAS-low Effector CD8+ T cells; (H) GO functional enrichment analysis of differentially expressed genes in the KRAS-high subset; (I) Immunofluorescence co-localization images in human carotid artery plaques.

Based on TCellSI scoring, the KRAS-high group displayed stronger cytotoxicity, exhaustion, and senescence phenotypes (Figure 5G). Genes upregulated in the KRAS-high group relative to the KRAS-low group were enriched in GO terms such as immune response, positive regulation of gene expression, and cell surface receptor signaling pathway, indicating that KRAS-high effector CD8+T cells are in a more active effector state within the plaque microenvironment (Figure 5H). Immunofluorescence co-localization staining of human advanced carotid atherosclerotic plaques further confirmed the presence of this subpopulation (Figure 5I).

These findings collectively suggest that KRAS-high CD8+T cells exhibit enhanced cytotoxic potential and tissue residency within atherosclerotic lesions, potentially driving local inflammation and plaque destabilization. Their enrichment correlates with increased IFN-γ and granzyme B expression, reinforcing their role in macrophage activation and vascular smooth muscle cell apoptosis.

### Deep learning model combined with single-cell transcriptomic sequencing predicts candidate therapeutic compounds

To identify small molecules capable of reversing the transcriptional abnormalities induced by KRAS upregulation in T cells of patients with premature CAD, we employed the deep learning model DrugReflector for compound prioritization. DrugReflector, trained on the Connectivity Map (CMap) covering 9,597 perturbations across 52 cell lines, ranks candidate compounds based on their predicted ability to reverse user-defined gene expression signatures^[18]^. We input the differentially expressed genes between T cells of the MI and Healthy groups obtained from our single-cell analysis as the disease signature (Figure 6A-D). The prediction results showed that BRD-K78432605 achieved the highest matching score, suggesting that this compound is most likely to bind KRAS or modulate its downstream pathways in T cells, thereby redirecting the disease-associated transcriptome toward a healthy state (Figure 6E and F).

**Figure 6.**
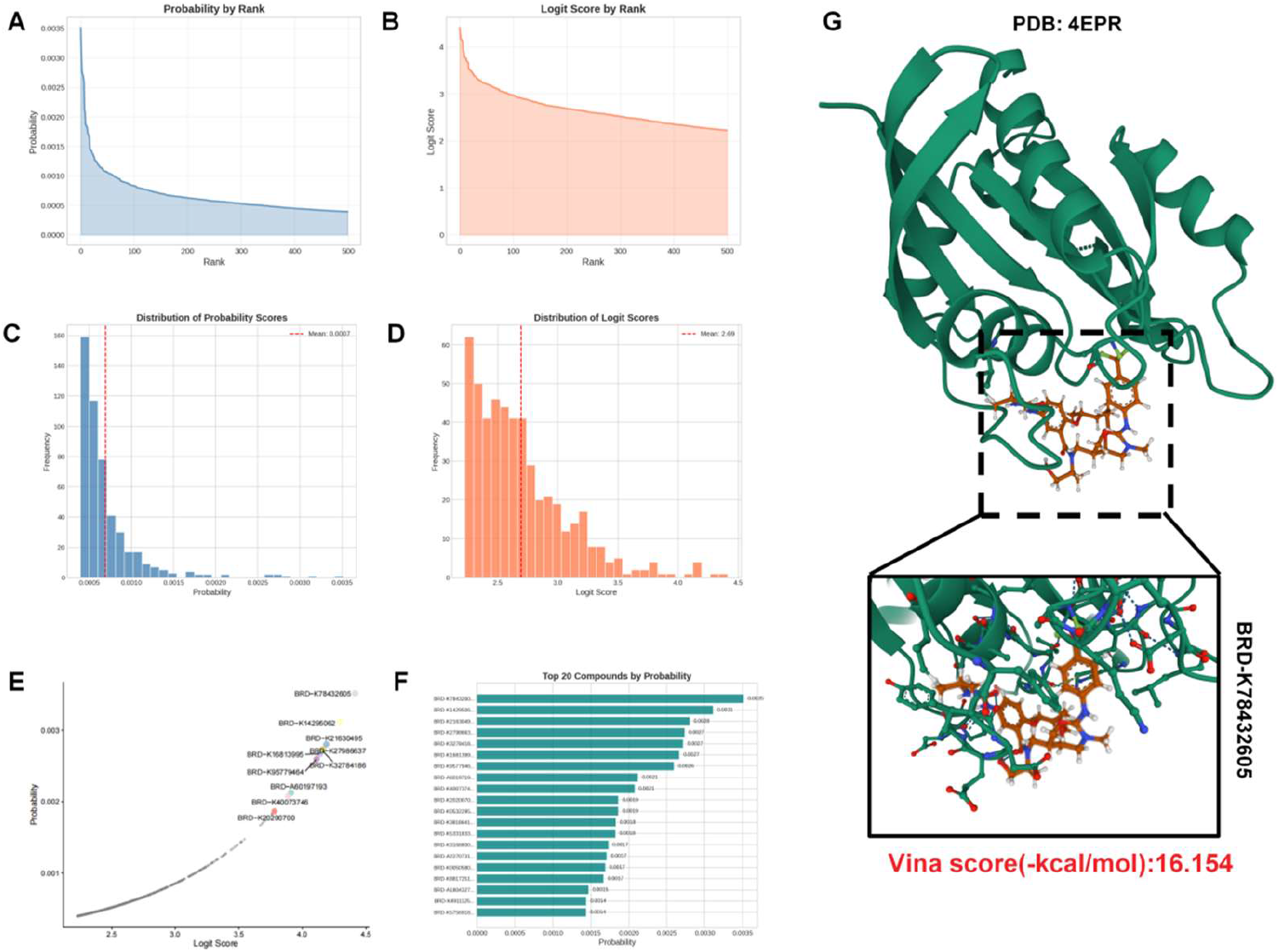
Drug prediction by the DrugReflector deep learning model and molecular docking validation. A–B) Probability and Logit Score (ranked) of compounds predicted by DrugReflector after inputting PBMC single-cell results; (C–D) Distribution plots of Probability and Logit Score of compounds predicted by DrugReflector after inputting PBMC single-cell results; (E–F) Top 10 and Top 20 compounds displayed in the Probability and Logit Score prediction curves; (G) Schematic diagram of the molecular docking conformation of BRD-K78432605 and the KRAS closed conformation (PDB: 4EPR).

To validate this prediction and preliminarily characterize its binding mode, we performed molecular docking to assess the affinity of BRD-K78432605 for different conformations of the KRAS protein. Because no high-resolution crystal structure of wild-type KRAS bound to GTP or its analog is currently available in the PDB database, we used the GDP-bound inactive closed conformation (PDB: 4EPR, resolution 2.00 Å, wild-type) as the docking receptor. The molecular docking result yielded a binding energy of −16.154 kcal/mol (Figure 6G). This value is substantially lower than the widely used activity threshold of −7.0 kcal/mol, indicating a thermodynamically favorable binding tendency, and suggests that BRD-K78432605 may form stable contacts at the SOS-binding interface of the closed conformation. These findings provide preliminary evidence that this small molecule has the potential to bind KRAS in its inactive state and to stabilize this conformation.

## DISCUSSION

By integrating transcriptomic profiling, single-cell transcriptomics, virtual gene knockout, deep learning-based prediction, and molecular docking, this study reveals the upregulation of the proto-oncogene KRAS in peripheral blood T cells of patients with PCAD, delineates its potential functional consequences, and validates the presence of this aberrant T-cell subpopulation within atherosclerotic plaques. This finding expands our understanding of KRAS function in the cardiovascular immune microenvironment.

We identified KRAS from the Aging Atlas database and observed, at the single-cell level, that effector CD8+T cells with high KRAS expression exhibit the highest senescence scores; these scores are further elevated in the MI group compared with healthy controls, indicating that these cells are undergoing accelerated immunosenescence. This observation offers an immunosenescence-centered perspective on PCAD pathogenesis: in young individuals, specific T-cell subsets may prematurely acquire a senescence-associated phenotype. The SASP factors they release, together with their heightened cytotoxicity, collectively create an immune microenvironment that promotes vascular injury. Notably, we found that overall cell–cell communication in PBMCs was diminished in the MI group relative to the healthy group, a pattern that mirrors the loss of intercellular coordination characteristic of the aging immune system, further supporting a link between PCAD and immunosenescence. Therefore, KRAS upregulation in T cells may represent a key molecular event connecting immunosenescence to premature atherosclerosis.

The identification and risk stratification of PCAD remain substantial clinical challenges^[19]^. The 2018 AHA/ACC multi-society cholesterol guideline recommends considering coronary artery calcium (CAC) scoring to guide treatment decisions, yet this recommendation rests primarily on data from the Multi-Ethnic Study of Atherosclerosis (MESA), which enrolled adults aged ≥45 years, and its extrapolation to younger adults has not been validated^[20]^. Likewise, the Pooled Cohort Equations used to estimate 10-year atherosclerotic cardiovascular disease (ASCVD) risk are not applicable to individuals younger than 40 years. Although the estimated 10-year risk in young adults may appear low, their lifetime risk is not necessarily negligible^[21]^. The proportion of acute myocardial infarction occurring in young individuals has risen in recent years; while these patients often present with less extensive coronary atherosclerosis and a higher prevalence of single-vessel disease, their long-term prognosis remains poor^[22]^.

Specific biomarkers that could aid in the diagnosis of PCAD have yet to be identified.In a comparative study of potential biomarkers, C-reactive protein was the only marker that remained significantly elevated in patients with PCAD after multivariate adjustment^[23]^. Another study confirmed the independent prognostic value of the plasma atherosclerosis index (AIP) for major adverse cardiovascular events in this population; however, it has not yet been incorporated into routine clinical practice^[24]^. Moreover, young patients with PCAD exhibit a higher burden of psychological distress and poorer medication adherence, further complicating management^[25]^. Our discovery of KRAS upregulation in peripheral blood T cells—linked to accelerated immunosenescence and cytotoxic effector programs—raises the possibility that KRAS or its downstream transcriptional signatures can serve as novel early-warning biomarkers that reflect the premature biological aging of the immune system in PCAD, offering a complement to traditional lipid-based and inflammatory markers. Prospective studies that correlate KRAS expression in circulating T-cell subsets with plaque progression and clinical events will be needed to determine whether this molecular feature can improve risk discrimination and guide earlier, more personalized intervention in young adults at risk.

We noted that CCR7 was the most significantly perturbed gene upon virtual knockout of KRAS. CCR7 is a key chemokine receptor that directs naïve T cell homing to secondary lymphoid organs; its ligands, CCL19 and CCL21, are produced by lymphoid stromal cells and guide T cells to efficiently scan antigen-presenting cells within lymphoid tissues^[26]^. However, a recent study published in Science revealed that CCR7 signaling extends well beyond its classical navigational function: after CD8+T cells form immunological synapses with dendritic cells and become activated, CCR7 signaling can serve as an early checkpoint that drives the active detachment of activated T cells from dendritic cells, thereby ensuring timely termination of contact with antigen-presenting cells. When CCR7 signaling is blocked or absent, CD8+T cell–dendritic cell interactions become abnormally prolonged, causing effector T cells to acquire a dysfunctional phenotype characterized by elevated expression of inhibitory receptors such as PD-1 and CTLA-4 and impaired antimicrobial activity^[27]^.

This finding has direct implications for understanding the behavior of KRAS-upregulated T cells in premature coronary artery disease. When KRAS is aberrantly upregulated, CCR7 expression may be persistently suppressed, depriving activated T cells of the detachment signal. This has two critical consequences: first, T cells prematurely escape homing surveillance in lymphoid organs and are more prone to migrate to and accumulate in peripheral tissues, including the arterial wall; second, owing to the loss of CCR7-mediated early checkpoint control, T cells within the plaque microenvironment may undergo sustained TCR signal integration, mirroring the failure to appropriately terminate activation observed when CCR7 signaling is disrupted, and ultimately acquire an exhausted-senescent phenotype marked by high expression of inhibitory receptors and dysfunctional effector activity. This aligns closely with our observation that effector CD8+T cells in plaques simultaneously exhibit enhanced cytotoxicity, elevated exhaustion scores, and increased senescence scores—an apparently paradoxical coexistence of high killing capacity, high exhaustion, and senescence that may fundamentally arise from the loss of CCR7-dependent regulation of activation kinetics driven by KRAS upregulation.

This study positions KRAS-upregulated senescent cytotoxic CD8+ T cells as a novel link between immunosenescence and premature atherosclerosis. Future work should prospectively evaluate KRAS signatures in young cohorts for risk stratification, establish causality using T-cell-specific Kras gain/loss-of-function models, and dissect CCR7-dependent checkpoint dysregulation as a potential therapeutic target.

## LIMITATIONS

Several limitations of this study should be acknowledged. First, owing to the absence of complete clinical follow-up information in the public datasets, we were unable to directly assess the association between KRAS expression levels in T cells and clinical outcomes such as major adverse cardiovascular events, which limits the prognostic interpretation of KRAS as a biomarker. Second, the datasets were derived from different disease contexts, including coronary artery disease in patients under 45 years of age, myocardial ischemia(ischemic cardiomyopathy), and advanced atherosclerotic plaques. Although these conditions collectively belong to the spectrum of premature atherosclerosis, their disease stages and clinical definitions are not fully equivalent, and combining them may introduce heterogeneity. Third, the single-cell analysis of PBMCs was based on a very limited number of donors (two disease samples and four controls), which may amplify individual variation; however, the key findings were corroborated in independent plaque scRNA-seq datasets and by immunofluorescence staining. Fourth, all mechanistic inferences in this study are based on computational analyses and re-analysis of existing sequencing data, without direct KRAS gain- or loss-of-function validation. Whether KRAS upregulation is sufficient to drive the acquisition of a cytotoxic, senescent T-cell phenotype remains to be established through experimental approaches such as lentiviral overexpression in human T-cell lines or primary T cells and conditional T-cell-specific Kras knock-in models. Fifth, we did not examine whether KRAS mutations are present in peripheral blood T cells of patients with premature coronary artery disease; future studies should include KRAS gene sequencing to clarify the relative contributions of transcriptional upregulation versus rare mutations. Finally, the candidate compound BRD-K78432605 identified by computational prediction requires experimental validation before any therapeutic relevance can be claimed.

## DECLARATIONS

## Acknowledgments

The authors gratefully acknowledge the Gene Expression Omnibus (GEO) database for providing the publicly available datasets used in this study. We also acknowledge colleagues and collaborators who offered valuable discussions and technical suggestions during the course of this research, as well as the biobank and clinical teams who facilitated tissue specimen acquisition and processing.

## Authors’ contributions

Zhao Sun, Ziyu Yang, and Xuedong Wang contributed equally to this work as co-first authors. They made substantial contributions to the conception and design of the study, performed bioinformatics analyses including bulk RNA-seq differential expression analysis, WGCNA, single-cell RNA-seq data processing, virtual gene knockout, and deep learning-based drug prediction, and were primarily responsible for data interpretation and manuscript drafting.

Meng Zhang and Rui Gao contributed to data acquisition, processing of public datasets, and provided technical support for single-cell analysis and molecular docking simulations.

Mengyue Yang and Yanchao Li performed the PCR validation experiments and immunofluorescence staining, and contributed to the acquisition and analysis of experimental data.

Qi Liu and Jingbo Hou jointly supervised the study, contributed to the conception and design of the research, provided funding and administrative support, and critically revised the manuscript for important intellectual content. Both are designated as corresponding authors.

All authors reviewed and approved the final version of the manuscript and agreed to be accountable for all aspects of the work.

## Availability of data and materials

The bulk RNA-seq (GSE12288) and scRNA-seq data (GSE231906, GSE145154, GSE252243 and GSE264666) used in this study were all obtained from the publicly available GEO database and are freely accessible.

## AI and AI-assisted tools Statement

The authors declare the use of artificial intelligence (AI) and AI-assisted technologies in the data analysis and prediction components of this study.

scTenifoldKnk (version 1.0.3) was used for in silico gene knockout analysis. This tool constructs single-cell gene regulatory networks, removes a target gene, and compares the resulting network with the original to identify significantly perturbed genes, thereby inferring the gene’s regulatory role.

DrugReflector was employed for transcriptome-driven drug prediction.

DrugReflector is a deep learning model trained on the Connectivity Map (CMap) database; it ranks small-molecule compounds based on their predicted ability to reverse a user-defined gene expression signature.

No generative AI or AI-assisted tools were used for data analysis or content generation in this work, except for grammar and spelling correction.

## Ethical approval and consent to participate

For the immunofluorescence staining experiments, formalin-fixed paraffin-embedded human carotid atherosclerotic plaque specimens were obtained from Department of Neurosurgery, The Second Affiliated Hospital of Harbin Medical University. The use of these archival specimens was approved by the Ethics Committee of The Second Affiliated Hospital of Harbin Medical University (approval number:KY-2025394), and written informed consent was obtained from all participants or their legal representatives prior to tissue collection.

## Consent for publication

Not applicable.

## Copyright

© The Author(s) 2023.

